# The architecture of the actin network can balance the pushing forces produced by growing microtubules

**DOI:** 10.1101/2022.01.21.476947

**Authors:** Shohei Yamamoto, Jérémie Gaillard, Benoit Vianay, Christophe Guerin, Magali Orhant-Prioux, Laurent Blanchoin, Manuel Théry

## Abstract

The position of centrosome, the main microtubule-organizing center (MTOC), is instrumental in the definition of cell polarity. It is defined by the balance of tension and pressure forces in the network of microtubules (MTs). As MTs polymerize against the cell periphery, pressure increases and produces pushing forces on the MTOC. How the mechanical interplay between MTs and the actin network is involved in the regulation of these forces remains poorly understood, in particular because its investigation is technically limited by the structural and biochemical complexity of the cell cytoplasm. Here, in a cell-free assay, we used purified proteins to reconstitute the interaction of an aster of dynamic MTs with actin networks of various compositions and architectures in cell-sized microwells. In the absence of actin filaments, the positioning of the MTOC was highly sensitive to variations in MT length. The presence of a bulk actin filament network limited MTs deformation and displacement, and MTOCs were hold in place. In contrast, the assembly of a dense and branched actin network along the edges of the wells centered the MTOCs by preventing MT slippage and thus maintaining an isotropic balance of pushing forces. In agreement with this, an asymmetric peripheral actin network caused the MTOC to decenter by creating an asymmetry in the pushing forces. Numerical simulations demonstrated that steric hindrance by actin networks, at the tip or along the entire length of MTs, can modulate MTOC positioning, as observed in the experiments. Overall, our results show that actin networks can limit the sensitivity of MTOC positioning to MT length and enforce robust MTOC centering or decentering depending on its architecture.

## Introduction

The network of microtubules (MTs) supports the construction of cell body plan and directs its symmetry axes ^1–3^. The asymmetric organization of MTs directs the preferential orientation of vesicle transport and the position of key sensory organelles and thereby orients the main functions of polarized cells ^2,4–6^. The centrosome is the main microtubule-organizing center (MTOC), so its position is a key determinant of cell polarity ^1,7^. The positioning of centrosome, either at the cell center or at the cell periphery, in contact with the plasma membrane, is important for ciliogenesis, immune reactions, cell division, epithelial-to-mesenchymal transition or neuronal development ^8–12^.

Centrosome position is mainly controlled by combinations of pushing and pulling forces produced in the MT network ^13–18^. Homogeneous distribution of minus-end directed molecular motors, pulling on MTs in the cytoplasm or at the cell cortex, can enforce centrosome centering ^19–22^. Heterogeneous distribution of motors, due to local accumulations, can locally increase pulling forces and enforce MTOC off-centering up to the contact with the plasma membrane ^23,24^. Regulation of centrosome positioning by such pulling forces is robust: centering, in the case of homogeneous distribution of motors, or peripheral positioning, in the case of heterogeneous distribution, are both poorly sensitive to variations of MT length ^13^. On the opposite, production of pushing forces is much more sensitive to variations of MT length and hence appears as a less reliable positioning mechanism. MT-based pushing is ineffective if MTs are too short to reach the spatial boundaries. Longer MTs may allow centering if their length corresponds precisely to the length of its confining region. However, MTs longer than this critical length will induce an abrupt transition from centering to off-centering ^20,25–27^.

Actin networks are involved in centrosome positioning and have been proposed to modulate the pushing forces produced by MTs but the underlying mechanisms are obscure ^14,18,28–31^. There are numerous examples of physical interactions between MTs and actin filaments ^32–37^. Acting as obstacles, capturing sites or stabilizing sheaths, actin architectures may regulate and organize the spatial distribution of forces in the MT network, by either amplifying or buffering local asymmetries. However, how specific actin architectures, such as dense cortical networks, bundles of linear filaments or cytoplasmic mesh, specifically impact forces production and propagation along MTs, remains poorly understood.

Here, we show that the presence and architecture of actin networks affect the spatial distribution of MTs and the displacements of MTOC either toward or away from the geometrical center of the well. Our results revealed how various actin networks have distinct and specific impact on the distribution of pushing forces and confer some robustness to the mechanism of MTOC positioning by making it less sensitive to variations of MT length.

## Results

### aMTOC positioning in microwells

It is difficult to directly assess the mechanical effects of actin networks on the production of pushing forces by MTs in living cells because of the biochemical and structural complexities of cell interior, and the presence of motors exerting pulling forces on MTs. Therefore, to reconstitute the interaction of an aster of dynamic MTs with actin networks, we designed an *in vitro* reconstitution assay using purified proteins and microfabrication techniques.

We investigated how an astral array of MTs self-organizes in a cell-sized compartment. 3D microwell can be used to impose physical barrier to MT growth ^20,38^. As compared to lipid droplets ^25,39^, microwells offer the possibility to control the size and shape of the container ^40^. Actual boundaries are made of lipids in living systems. Therefore, we started by setting up the coating of microwell with a lipid bilayer ^41^ and further closed the upper surface of the container with a layer of mineral oil (Figure 1A-B, Supplementary Figure 1A-B). Lipids were properly diffusing in the bottom plane and the vertical edges of the microwell (Supplementary Figure 1C). However, there was a problem of tubulin precipitation, as reported previously ^42,43^. After 30 to 60 minutes of incubation of tubulin in the TicTac or the BRB80 buffer, the tubulin precipitated (Supplementary Figure 2A) and microtubule dynamics stopped. A screening of the biochemical conditions (Supplementary Figure 2B-F) suggested that at lower temperature (22°C) and with high concentration of BSA and GTP, tubulin precipitation could be delayed, up to two hours (Figure 1C). As purified centrosomes generate a quite variable number of MTs ^33^, we chose to work with artificial MTOCs (aMTOCs) made of short stabilized pieces of MTs grafted on a polystyrene bead (Figure 1D). They efficiently generated 15 to 20 dynamic MTs per aMTOC (Supplementary Figure 3A-D).

**Figure 1.**
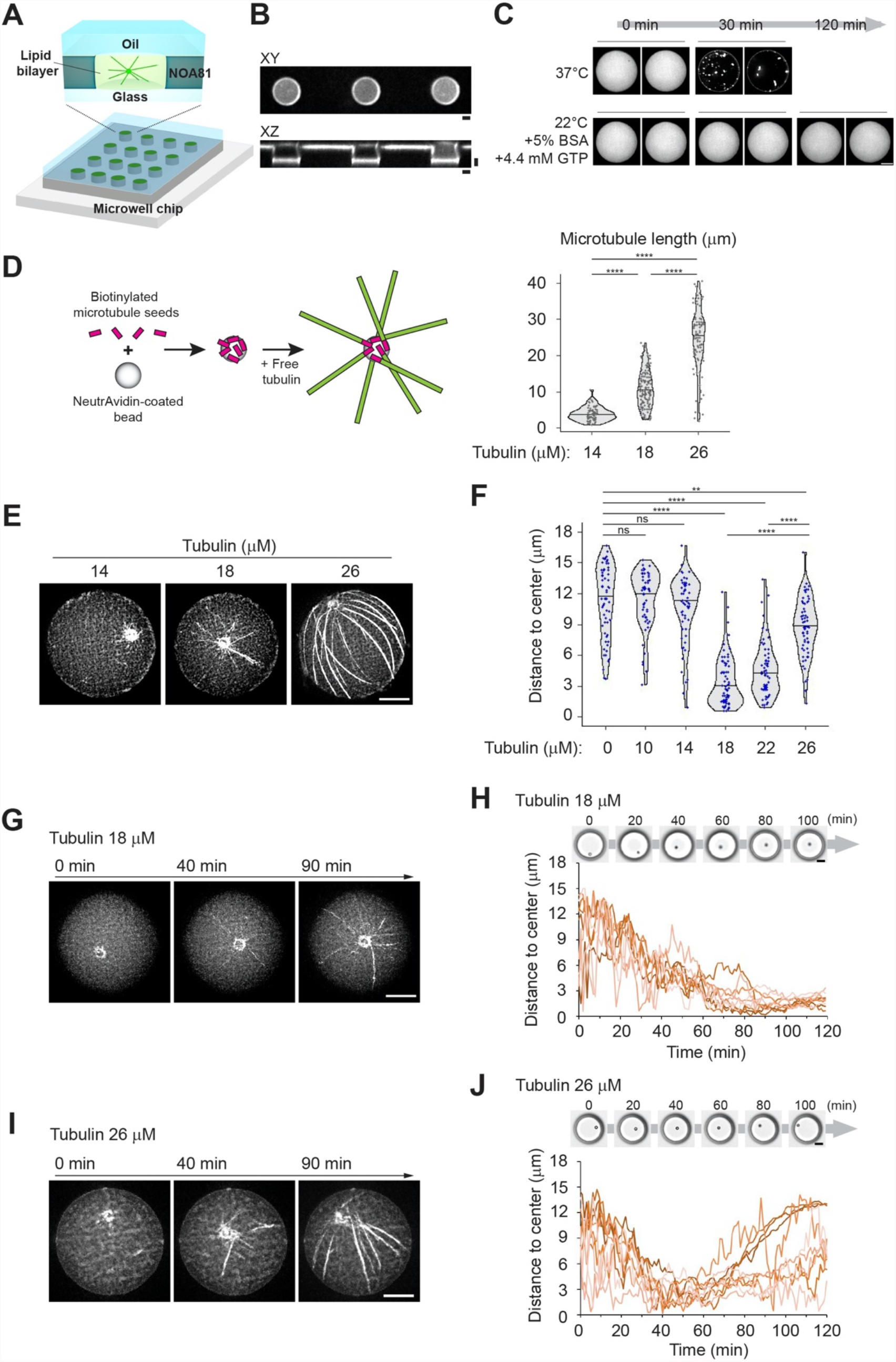
aMTOC positioning in microwells. **(A)** Scheme of microwells (See also Supplementary Figure 1A, B). **(B)** Images of fluorescence-labelled lipid. XY and XZ view of microwells were shown. **(C)** A screening of biochemical conditions to slow down tubulin precipitation (See also Supplementary Figure 2 and Methods). **(D)** Scheme of preparation of an artificial MTOC (aMTOC). Biotinylated MT seeds were attached on NeutrAvidin-coated beads. By the addition of free tubulin, MT polymerization occurs from the beads (See also Methods). Right graph shows MT length at the indicated tubulin concentrations in microwells. (Tubulin 14 μM n = 71, 18 μM n = 134, 26 μM n = 97 MTs (8 wells, respectively)) ****p<0.0001 (Kruskal-Wallis test with Dunn’s multiple comparison test). **(E)** Representative images of MT asters with various tubulin concentrations. Images were taken at 2 hours after sample preparation. **(F)** Distance from aMTOC to center of the well at the indicated tubulin concentrations (2 hours after sample preparation). (Tubulin 0 μM n = 62, 10 μM n = 59, 14 μM n = 60, 18 μM n = 68, 22 μM n = 60, 26 μM n =65 wells) **p<0.01, ****p<0.0001, ns (not significant)>0.1 (Kruskal-Wallis test with Dunn’s multiple comparison test). **(G)** Time-lapse imaging of MT aster formation at 18 μM of tubulin. **(H)** aMTOC position over time at 18 μM of tubulin. Bright-field images were taken at 1 min intervals. Positions of 10 individual aMTOCs were shown with different colors. **(I)** Time-lapse imaging of MT aster formation at 26 μM of tubulin. **(J)** aMTOC position over time at 26 μM of tubulin. Bright-field images were taken at 1 min intervals. Positions of 10 individual aMTOCs were shown with different colors. Scale bar 10 μm. Violin plots were shown with the median (horizontal line).

We first analyzed the sensitivity of aMTOC positioning to the ratio of MT length over container length by varying tubulin concentrations in microwells of controlled size. As tubulin concentration has been increased from 14 to 26 μM, the average length of MTs varied from 4 to 25 μm (Figure 1D, E). The radius of the microwell was close to 19 μm. Below 18 μM, MTs were shorter than 10 μm and aMTOC adopted a random position (Figure 1F and Supplementary Figure 4A, B). At 18 μM of tubulin, MTs were longer and could reach the microwell boundaries (Figure 1D, E). In about 60 min (Figure 1G, H and Supplementary Movie 1), most aMTOC reached the center of the microwell and remained there (Figure 1F and Supplementary Figure 4B). Centering was also efficient at 22 μM of tubulin (Figure 1F). At 26 μM of tubulin, most MTs were longer than 20 μm (Figure 1D). As they grew, they first ensured a proper centering but after an hour, MT elongation and slippage along microwell edges broke the network symmetry and MTs pushed aMTOC away from the center (Figure 1I, J and Supplementary Movie 2). Taken together, in our experimental system, aMTOC positioning appeared highly sensitive to the tubulin concentration and the MT length. Above a critical concentration of 22 μM, MT elongation and reorientation could bias the distribution of pushing forces and promote aMTOC off-centering, as previously described in water-in-oil droplets ^25^.

Previous works based on numerical simulations suggested that the friction along the boundaries of the container might prevent the symmetry break by enforcing a vortex-like structure in the network that preserves MTOC centering ^13^. In cells, the actin filaments form distinct cortical and cytoplasmic networks that might restrict MT lateral translocation and aster displacement ^44,45^. Therefore, we tested how these various architectures might impact either centering or off-centering mechanisms.

### Assembly of various actin architectures in microwells

Rather than working with preassembled and stabilized actin filaments, we chose to grow actin filaments in the microwells in order to control their position and architecture by controlling the mechanism of actin network assembly. Unbranched actin network could be formed in the bulk of the microwell simply by spontaneously assembling 4 μM of actin monomers (Figure 2A, B and Supplementary Figure 5A). Alternatively, to limit the cytoplasmic pool, and favor the assembly of a dense cortical layer, actin filaments were nucleated near the lipid layer by a Nucleation Promoting Factor (NPF) attached to the lipid in the presence of the Arp2/3 complex and actin monomers in the solution (Figure 2C, D and Supplementary Figure 5A, B).

**Figure 2.**
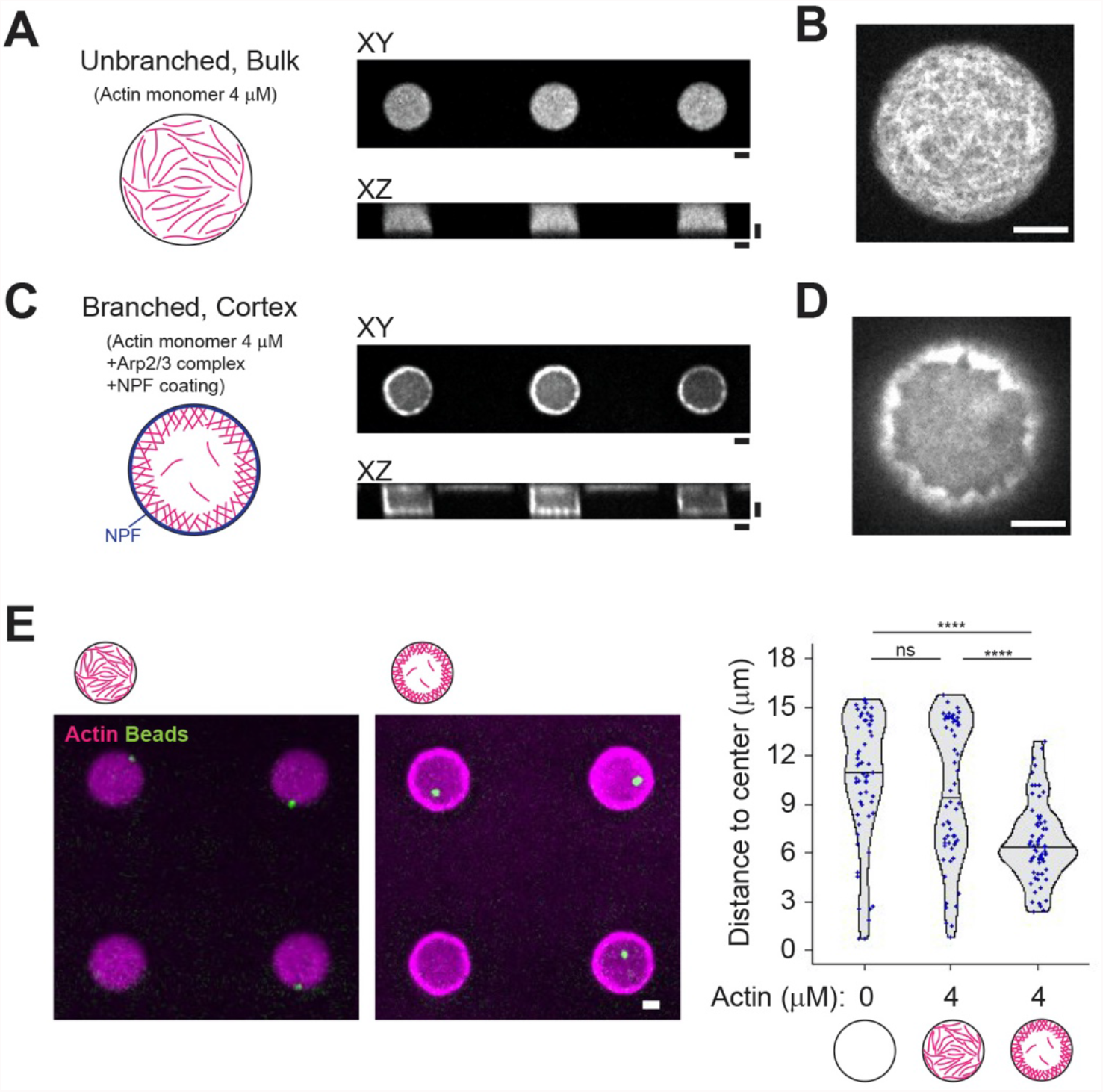
Assembly of various actin architectures in microwells. **(A)** and **(B)** Unbranched, bulk actin network. XY and XZ views were shown. Higher magnification image (XY view) was shown in **(B). (C)** and **(D)** Cortical branched actin network. XY and XZ views were shown. NPF (streptavidin-tagged WA) was coated on the lipid-biotin. Higher magnification image (XY view) was shown in **(D). (E)** aMTOC position in the absence of free tubulin. Left, representative images showing actin and the aMTOCs in microwells. Right, measurement of distance from aMTOC to center of the well (2 hours after sample preparation). Violin plots were shown with the median (horizontal line). (Actin 0 μM n = 60, Actin 4 μM Bulk n = 60, Actin 4 μM Cortex n = 63 wells) ****p<0.0001, ns (not significant)>0.1 (Kruskal-Wallis test with Dunn’s multiple comparison test). Scale bar 10 μm.

We first tested the impact of these two actin architectures on aMTOC positioning independently of MTs (Figure 2E). Unbranched actin filaments in the bulk had no visible impact on bead position as compared to similar conditions without them (Figure 2E and Supplementary Figure 5C). Position of the beads was a bit centered in the presence of cortical actin, as the thickness of the cortical layer restricted the available space for beads (Figure 2E and Supplementary Figure 5C, D).

These results showed that actin filaments in the bulk and branched cortical network can be reconstituted in 3D microwells and have distinct impact on aMTOC position independently of MTs. The two networks may also have specific effects on MT slippage or deformation and as such distinct impacts on the force distribution in the MT network and therefore on the positioning of the MTOC.

### Bulk actin network impairs aMTOC displacement and aster self-centering

aMTOC displacements depend on the production of pushing forces by MT polymerization against container boundaries. However, the morphology of aster confers them a large effective cross-section that limits their displacements by viscous drag. We first tested whether the density of actin network in the bulk could impact the production of pushing forces against effective boundaries, reorganize the spatial distribution of MTs and thus affect the inner balance of force production by MT polymerization.

To test whether bulk actin network could impair the centering process, we worked in conditions where MT length was comparable to the microwell radius (ie at 18μM tubulin). aMTOC position was not dramatically affected by 1 μM of actin but appeared off-centered with 4 μM of actin (Figure 3A, B-left and Supplementary Figure 6A). Indeed, time-lapse imaging revealed that aMTOC remained stuck at their initial position in the presence of 4 μM of actin (Figure 3C). Importantly, at this concentration, unbranched actin network had no impact on MT elongation (Figure 3D). In addition, higher concentrations of tubulin, 26 μM, although capable of promoting MT elongation, could not overcome aster immobilization by the bulk actin meshwork (Figure 3B-right and Supplementary Figure 6B). This suggested that the defective centering was due to friction resisting aster displacement rather than steric effects blocking MT polymerization ^32,33^. Indeed, implementing physical hindrance to aster displacement by taking into account steric effect of actin filaments along MT aster in numerical simulations was sufficient to account for the immobilization of MTOC by linear bulk actin filaments (Figure 3E-G, Supplementary Figure 6C, D and Supplementary Movie 3). Therefore, we conclude that the presence of a dense network of actin filaments in the bulk can affect MT aster centering by resisting aster translocation rather than impairing MT elongation.

**Figure 3.**
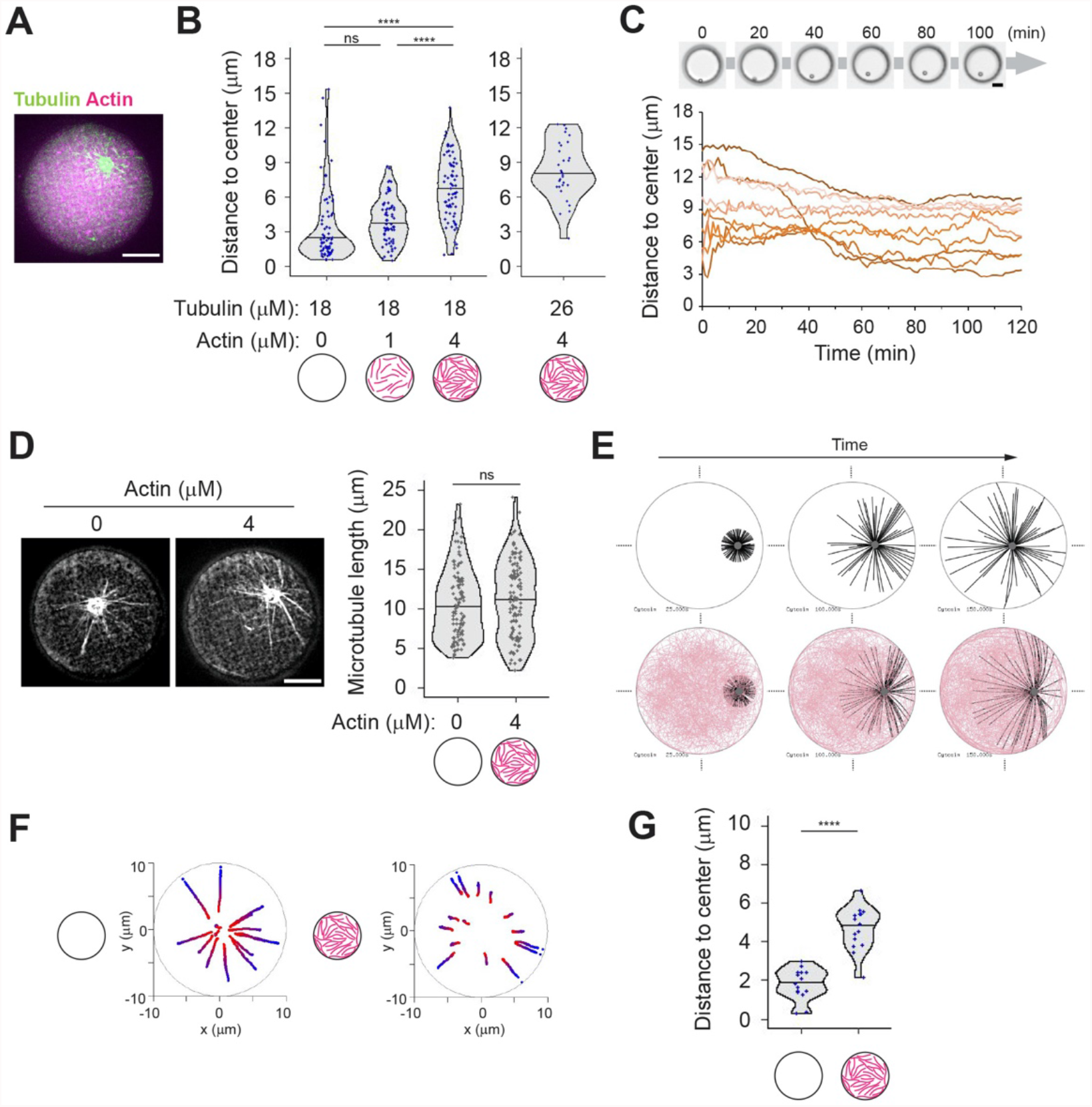
Bulk actin network impairs aMTOC displacement and aster self-centering. **(A)** Representative image of MT aster in the presence of bulk actin network. Tubulin 18 μM and actin 4 μM. **(B)** Distance from aMTOC to well center (2 hours after sample preparation). Left: Tubulin 18 μM. (Actin 0 μM n = 68, 1 μM n = 65, 4 μM n = 71 wells). Right: Tubulin 26 μM Actin 4 μM (n = 30 wells). ****p<0.0001, ns (not significant)>0.1 (Kruskal-Wallis test with Dunn’s multiple comparison test). **(C)** aMTOC position over time. Tubulin 18 μM and Actin 4 μM. Bright-field images were taken at 1 min intervals. Positions of 10 individual aMTOC were shown with different colors. **(D)** Measurement of MT length in the absence or presence of bulk actin network. Tubulin 18 μM. Images were taken 2 hours after sample preparation. (Actin 0 μM n = 109, 4 μM n = 118 MTs (from 8 wells, respectively)). ns (not significant)>0.1 (Mann-Whitney U test). **(E)** Simulations in the absence (top) or presence (bottom) of actin filaments. Different time points (From left, 25, 100 and 150 sec) were shown. MTOC, gray, MT, black, Actin, pink. **(F)** Trajectories of MTOCs from blue (0 sec) to red (150 sec). 15 simulations per condition. The initial position (0 sec) was randomly chosen. **(G)** Final position of MTOC (at 150 sec). 15 simulations per condition. ****p<0.0001 (Mann-Whitney U test). Violin plots were shown with the median (horizontal line). Scale bar 10 μm in (A), (C) and (D).

These data also suggested that restricting the actin network to the periphery might specifically limit MT slippage without impairing aster translocation.

### Cortical branched actin meshwork favors aster centering

To test whether cortical actin could counteract MT slippage and MTOC off-centering, experiments were performed in the presence of long MTs (ie at 26μM tubulin) (Figure 4A). As described above, the cortical network clustered aMTOC in a smaller volume and thus induced a partial centering (Figure 2E-right, 4B-left). Interestingly, this centering was significantly improved by the growth of long MTs (Figure 4B-left and Supplementary Figure 7A). MT length was not significantly changed even in the absence or presence of cortical actin, suggesting that this effect is not due to interference of actin with MT elongation (Figure 4C). Instead, MTs appeared longer than the radius of the microwell and the network adopted a vortex-like structure (Figure 4D). Time-lapse imaging showed that in the absence of cortical actin, MTs were pivoting around the aMTOC, whereas they maintained their orientation and grew along the edge in the presence of cortical actin (Figure 4E, F and Supplementary Movie 4). MT pivoting in the absence of cortical actin appeared associated with aMTOC off-centering, whereas MT sneaking into the cortical actin was associated with aMTOC stable centering and maintenance at the center (Figure 4G and H). Numerical simulations in which friction was restricted to the cell periphery displayed similar aMTOC displacements and final positions (Figure 4I-K, Supplementary Figure 7B and Supplementary Movie 5), demonstrating that local steric interactions between cortical actin and MTs are indeed sufficient to prevent MTOC off-centering. From these results, we concluded that cortical actin network can counteract the effect of MT elongation and aster off-centering by restricting MT slippage and thus maintaining a regular distribution of force application sites along the cortex and around the aMTOC. In addition, even at lower concentrations of tubulin (18 μM), MT asters displayed a proper centering mechanism in the presence of cortical actin (Figure 4B-right, and Supplementary Figure 7C). This showed that cortical actin can enforce a robust centering that is less sensitive to MT length.

**Figure 4.**
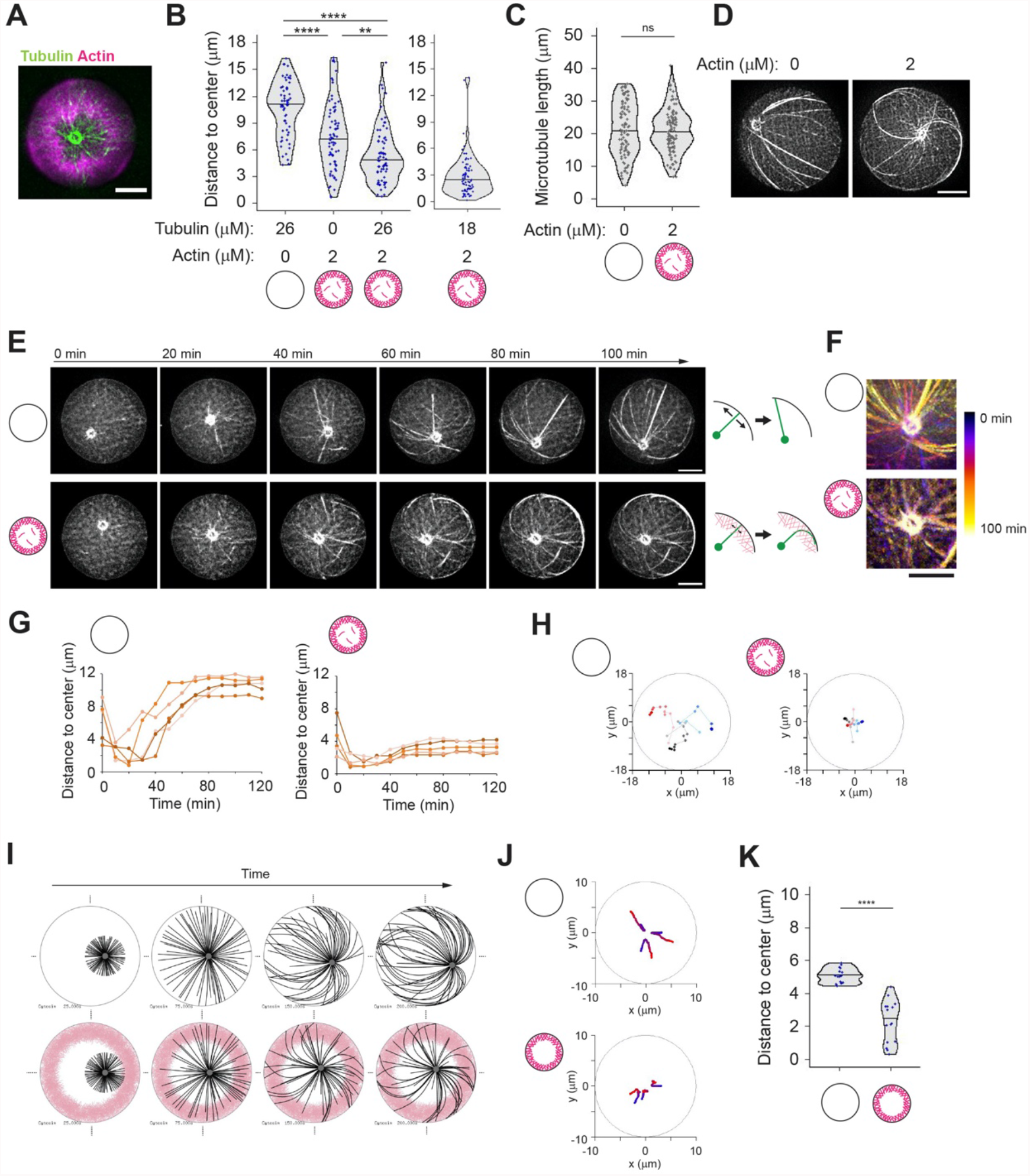
Cortical branched actin meshwork favors aster centering. **(A)** Representative image of MT aster with cortical actin. Partial maximum projection was shown. Tubulin 26 μM, actin 2 μM and Arp2/3 complex were added into NPF (WA) coated microwells. **(B)** Distance from aMTOC to well center (2 hours after sample preparation). Left: Tubulin 26 μM Actin 0 μM n = 61, Tubulin 0 μM Actin 2 μM n = 73, Tubulin 26 μM Actin 2 μM n = 70 wells. Tubulin 18 μM Actin 2 μM n = 76 wells. **p<0.01, ****p<0.0001, ns (not significant)>0.1 (Kruskal-Wallis test with Dunn’s multiple comparison test). **(C)** Measurement of MT length in the absence or presence of cortical actin. Tubulin 26 μM. Images were taken 2 hours after sample preparation. (Actin 0 μM n = 96, 2 μM cortex n = 104 MTs (from 6 wells, respectively). ns (not significant)>0.1 (Mann-Whitney U test). **(D)** Representative images of MT organization in the absence or presence of cortical actin. Tubulin 26 μM. **(E)** Time-lapse imaging of MT aster positioning in the absence (top) of presence (bottom) of cortical actin. Final actin structure was shown in (A). Right schemes indicate how MTs behave along cell boundary. In the absence of actin, MTs slipped and reoriented along well boundary as they grew. In contrast, MT reorientation was restricted in the presence of actin, although MTs can grow through actin network and along well boundary. **(F)** MT motion around the aMTOC shown in Figure 4E. Temporal-color coded images were shown. The position of aMTOC was centered at each time point in the image. **(G)** aMTOC position overtime in the absence (left) or presence (right) of cortical actin. 5 representative data per condition were shown. **(H)** Representative trajectories of aMTOCs in microwells from light colors (0 min) to dark colors (120 min). Time-lapse imaging was performed for 2 hours at 10 min intervals. 3 trajectories per condition were shown with different colors. **(I)** Simulations in the absence or presence of actin. Different time points (From left, 25, 75, 150 and 200 sec) were shown. MTOC, gray, MT, black, Actin, pink. **(J)** Representative trajectories of MTOC from blue (0 sec) to red (200 sec). The initial position (0 sec) was randomly chosen within 4 μm from the cell center. 3 simulations per condition. **(K)** Final position of MTOC (at 200 sec). 15 simulations per condition. ****p<0.0001 (Mann-Whitney U test). Violin plots were shown with the median (horizontal line). Scale bar 10 μm in (A) and (D)-(F).

These results also suggested that a heterogeneous pattern of cortical friction might create an asymmetry in the angular distributions of MTs, and an alignment of pushing forces leading to aMTOC off-centering.

### Asymmetric cortical actin meshwork induces aster off-centering

We reasoned that with lower actin filament density, which would be crosslinked to each other, we could enforce the asymmetry of the actin network growing from the walls of the microwells ^46^. Indeed, we found that with 0.5 μM instead of 2 μM of actin and 100 nM of α-actinin, the cortical network grew from all edges but formed an asymmetrically positioned ring (Figure 5A and Supplementary Figure 8A, B). The growth of MTs from the asters did not seem to have an impact on the asymmetric architecture of the actin network (Figure 5B). To analyze aster positioning with respect to this asymmetry, images were reoriented in order to align horizontally the center of the actin inner zone and the center of the microwell (Figure 5C, D and Supplementary Figure 8C, D). To test the potential guiding effect of asymmetric cortical actin networks, we worked in off-centering conditions, ie 26 μM tubulin, in which aMTOCs are randomly distributed in the microwell in the absence of actin (Figure 5E and Supplementary Figure 8E, F). By its thickness, the cortical actin network constrained asters positioning and limits aMTOC dispersion even in the absence of MTs (Figure 5F and Supplementary Figure 8E, F). However, as anticipated, the reorientation of the microwells with respect to the asymmetry of the actin network revealed that as MTs grew from the asters they shifted the aMTOCs toward the center of the actin inner zone (Figure 5G, H, I, Supplementary Figure 8E, F and Supplementary Movie 6). Interestingly, in conditions imposing shorter MTs and an efficient centering of the aster, ie 18μM of tubulin, the asters in the presence of the asymmetric actin network appeared off-centered toward the center of the actin inner zone as well (Figure 5J and Supplementary Figure 8G). Numerical simulations confirmed that a heterogeneous friction pattern due to variable thickness in the cortical actin network along microwell boundary was sufficient to push the MTOC away from the thicker actin layer and thus promote aster off-centering (Figure 5K-M, Supplementary Figure 8H and Supplementary Movie 7). Overall, these results showed that the cortical actin network architecture can direct the position of MTOCs, either at the center or away from it, depending on its heterogeneity. It controls the force balance at the MTOC by modulating the pattern of friction resisting MT slippage and thus directing the localization of the sites of application of pushing forces.

**Figure 5.**
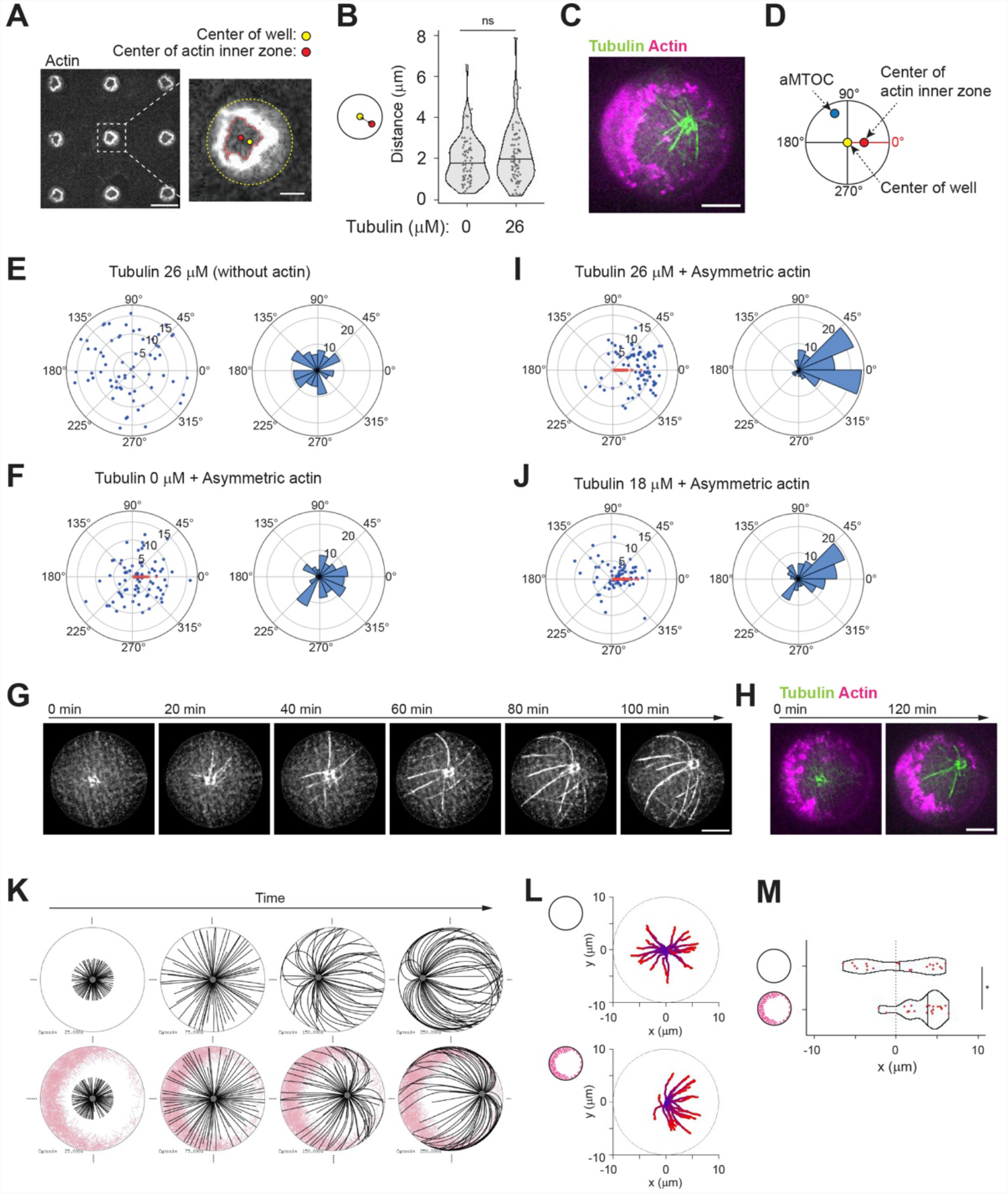
Asymmetric cortical actin meshwork induces aster off-centering. **(A)** Representative image of asymmetric actin cortex. Actin 0.5 μM, α-actinin 100 nM and Arp2/3 complex were added into NPF (WA) coated microwells. In the right image, the center of the well and the center of the actin inner zone were indicated. A single slice of the image was shown. Scale bar left 50 μm, right 10 μm. **(B)** Distance between well center and center of the actin inner zone. (Tubulin 0 μM n = 74, 26 μM n = 79 wells) ns (not significant)>0.1 (Mann-Whitney U test). **(C)** Representative image of MT aster with asymmetric actin. Tubulin 26 μM. Partial maximum projection was shown. **(D)** Scheme of angle measurements. Angles from the well center to the aMTOC and the center of the actin inner zone were measured. In (F), (I) and (J), the wells were reoriented based on each XY coordinates in order to align the angles from well center to center of the actin inner zone at 0° (See also Supplementary Figure 8C-D). **(E), (F), (I) and (J)** Distributions of aMTOCs. Left, aMTOC positions (μm) relative to well center. Right, Angular distributions (%) of aMTOCs from well center. (Tubulin 26 μM Actin 0 μM n = 65, Tubulin 0 μM Actin 0.5 μM n = 74, Tubulin 26 μM Actin 0.5 μM n = 79 wells, Tubulin 18 μM Actin 0.5 μM n = 68 wells,) Blue and red dots indicate the position of aMTOC and the center of the actin inner zone after alignment, respectively. **(G)** Time-lapse imaging of MT aster in the presence of asymmetric actin. Tubulin 26 μM. **(H)** Actin network structure of (H) at the initial (0 min) and final time point (120 min). Partial maximum projection was shown. **(K)** Simulations in the absence (top) or presence (bottom) of asymmetric actin (See also Supplementary Figure 8H). Initial position was set to the cell center. Time point: 25, 75, 150 and 250 sec. MTOC, gray, MT, black, Actin, pink. **(L)** Trajectories of MTOCs from blue (0 sec) to red (250 sec). 20 simulations. Initial position: cell center. **(M)** Final position of MTOC along X-axis (at 250 sec). 0 indicates the center along X axis in cells. 20 simulations per condition. *p<0.1 (Mann-Whitney U test). Violin plots were shown with the median (horizontal line). Scale bar, 10 μm in (C), (G) and (H).

## Discussion

Our results suggest that actin networks make aster positioning both more robust and more versatile. Indeed, in the absence of actin filaments, asters displayed abrupt transitions from centering to off-centering depending on MTs length. By contrast, bulk actin filaments resisted asters displacements and MTOC were hold in place, independently of the presence of MTs. Moreover, cortical actin networks specifically favored aster centering over a broad range of MT lengths. The presence of asymmetric actin resulted in off-centering of the aster. From these observations, we propose that actin networks can modulate the sensitivity of MT aster positioning to variation of MT length.

It has been reported that the presence of cytoplasmic actin network can affect MT organization ^45,47^. The immobilization of MTOC by bulk actin filaments in our system is reminiscent of the regulation of MT aster in Xenopus extract and sea urchin embryo ^32,48^. The robust polarization of MT network and off-centering of MTOC by asymmetric organization of the cortical actin is reminiscent of the mechanism supporting centrosome displacement to cell periphery during ciliogenesis and immune synapse formation ^8,11,49^. Thus, our reconstitution assay based on a minimal set of components might properly account for the actual mechanism regulating MT aster positioning by pushing forces in living cells. However, our experimental conditions and the use of short pieces of MTs attached to a bead did not offer us the possibility to control MT pivoting around the aMTOC, a property that could have promoted symmetry break and amplified MTOC off-centering ^13,50,51^.

Building a synthetic cell from scratch is a powerful strategy to improve our understanding of cell biology, and pave the way toward new living materials ^52^. Here, we established a way to combine dynamic MTs and actin filaments in cell-sized confinement. Our findings and techniques could be used as a step toward the reconstitution of the polarization process in synthetic cells.

## Acknowledgements

This work was supported by the European Research Council (Consolidator Grant 771599 (ICEBERG) to MT and Advanced Grant 741773 (AAA) to LB). S.Y. was supported by fellowships from the EMBO, Astellas Foundation for research on metabolic disorders and Mochida memorial foundation for medical and pharmaceutical research. This project was supported by the MuLife imaging facility, which is funded by GRAL, a program from the Chemistry Biology Health Graduate School of University Grenoble Alpes (ANR-17-EURE-0003).

## Materials and Methods

### Protein expression and purification

Tubulin was purified from fresh bovine brain by three cycles of temperature-dependent assembly/disassembly in Brinkley Buffer 80 (BRB80: 80 mM Pipes pH 6.8, 1 mM EGTA and 1 mM MgCl_2_) ^53^. MAP-free neurotubulin was purified by cation-exchange chromatography (EMD SO, 650 M, Merck) in 50 mM Pipes, pH 6.8, supplemented with 0.2 mM MgC1_2_, and 1 mM EGTA. Fluorescently labelled tubulin (ATTO-488- or ATTO-647-labelled) and biotinylated tubulin were prepared by following previously published method ^54^. Actin was purified from rabbit skeletal-muscle acetone powder. Monomeric Ca-ATP-actin was purified by gel-filtration chromatography on Sephacryl S-300 at 4°C in G buffer (2 mM Tris–HCl, pH 8.0, 0.2 mM ATP, 0.1 mM CaCl_2_, 1 mM NaN_3_ and 0.5 mM dithiothreitol (DTT)). Actin was labelled on lysines with Alexa-568. The Arp2/3 complex, recombinant GST-α-actinin 4 and GST-WA (a truncated version of human WASP) were purified in accordance with previous methods ^55,56^. Snap-Streptavidin-WA (pETplasmid) was expressed in Rosettas 2 (DE3) pLysS (Merck, 71403). Culture was grown in TB medium supplemented with 30 μg/mL kanamycine and 34 μg/mL chloramphenicol, then 0.5 mM isopropyl β-D-1-thiogalactopyranoside (IPTG) was added and protein was expressed overnight at 16 °C. Pelleted cells were resuspended in Lysis buffer (20 mM Tris pH8, 500 mM NaCl, 1 mM EDTA, 15 mM Imidazole, 0,1% TritonX100, 5% Glycerol, 1 mM DTT). Following sonication and centrifugation, the clarified extract was loaded on a Ni Sepharose high performance column (GE Healthcare Life Sciences, ref 17526802). Resin was washed with Wash buffer (20 mM Tris pH8, 500 mM NaCl, 1 mM EDTA, 30 mM Imidazole, 1 mM DTT). Protein was eluted with Elution buffer (20 mM Tris pH8, 500 mM NaCl, 1 mM EDTA, 300 mM Imidazole, 1 mM DTT). Purified protein was dialyzed overnight 4°C with storage buffer (20 mM Tris pH8, 150 mM NaCl, 1 mM EDTA, 1 mM DTT), concentrated with Amicon 3KD (Merck, ref UFC900324). Aliquots were flash frozen in liquid nitrogen and stored at −80 °C.

### Preparation of an artificial MTOC

To prepare microtubule seeds, the mixture containing 3 μM of fluorescent-labeled tubulin, 7 μM of biotinylated tubulin and 0.5 mM GMPCPP (Jena Bioscience, NU-405S) in BRB80 buffer was incubated at 37°C for 40 min. After the incubation, 10 μM of Taxol was added and the mixture was incubated at room temperature for 10 min. The microtubule seeds were then pelleted by centrifugation at 20,238 x g for 10 min and were resuspended in the BRB80 supplemented with 0.5 mM GMPCPP and 10 μM Taxol. The seeds were flash frozen and stored in liquid nitrogen. To prepare the Neutravidin-coated beads, the polystyrene beads containing surface primary amino groups (PolySciences, 17145-5, Diameter 3 μm) were incubated with 10 mM of Sulfo-NHS-LC-LC-Biotin (ThermoFisher, 21338) at room temperature for 40 min to modify their surface with biotin. The beads were washed with phosphate buffered saline (PBS) and then with HKEM buffer (10 mM HEPES (pH 7.5), 50 mM KCl, 5 mM MgCl_2_, 1 mM EGTA) supplemented with 0.1% bovine serum albumin (BSA). The beads were incubated with 1 mg/mL of Neutravidin (ThermoFisher, 31000) at 15°C for 30 min or at 4°C for 2 hours. After washing the beads with HKEM buffer supplemented with 0.1% BSA, the beads were resuspended in 200 μL of HKEM buffer supplemented with 0.1% BSA. The beads solution was then mixed with 10 μL of the microtubule seeds. The mixture was incubated under rotation at room temperature overnight. Before mixing the aMTOCs (microtubule seeds + beads) with the reaction mixture containing free tubulin, the solution containing aMTOCs was washed with HKEM supplemented with 0.1% BSA to remove excess seeds and Taxol.

### Preparation of small unilamellar vesicles (SUV)

L-α-phosphatidylcholine (EggPC) (Avanti, 840051C), 1,2-distearoyl-sn-glycero-3 phosphoethanolamine-N-[biotinyl(polyethylene glycol)-2000] (DSPE-PEG(2000)-Biotin) (Avanti, 880129C) and ATTO 647N labeled DOPE (ATTO-TEC, AD 647N-161 dehydrated) were used. Lipids were mixed in glass tubes as follows: Type 1 (99% EggPC (10 mg/mL) and 1% DOPE-ATTO390 (1 mg/mL)), Type 2 (98.75% EggPC (10 mg/mL) and 0.25% DSPE-PEG-Biotin (10 mg/mL) and 1% DOPE-ATTO390 (1 mg/mL)), Type 3 (99.5% EggPC (10 mg/mL), 0.25% DSPE-PEG-Biotin (10 mg/mL) and 0.25% DOPE-ATTO647N (1 mg/mL)). The mixture was dried with nitrogen gas. The dried lipids were incubated in a vacuum overnight. After that, the lipids were hydrated in the SUV buffer (10 mM Tris (pH 7.4), 150 mM NaCl, 2 mM CaCl_2_). The mixture was sonicated on ice. The mixture was then centrifuged for 10 min at 20,238 x g to remove large structures. The supernatants were collected and stored at 4°C. The final concentration of lipids was adjusted to 0.5 mg/mL. The Type 2 SUV was used to bind snap-streptavidin-WA onto the lipid layer. The Type 3 SUV was used to visualize lipids on microwells. In other experiments, the Type 1 SUV was used.

### Construction of microwells

The master mold (approximately 20 μm of thickness) was fabricated through photolithography using SU8 3025 (MicroChem) and then it was silanized by incubating with Trichloro (1H,1H,2H,2H-perfluoro-octyl) silane (Sigma, 448931). To make 1st PDMS, the mixture of prepolymer and curing agent (Dow, SYLGARD 184 silicone elastomer kit) was poured onto the master mold. It was baked at 70°C for 2 hours. The 1st PDMS was silanized by incubating with Trichloro (1H,1H,2H,2H-perfluoro-octyl) silane. The 2nd PDMS was made using the silanized 1st PDMS as a template. The 2nd PDMS was cut into small pieces and used as PDMS stamps.

Glasses were cleaned by successive chemical treatments: 30 min in acetone with sonication, 15 min in ethanol (96%), washing ultrapure water, 2 hours in HellmanexIII (2% in water, Hellma), washing in ultrapure water. The glasses were then dried. The slide glasses were oxidized in plasma cleaner (diener) for 2 min at power of 80% and then incubated overnight in a solution of 1 mg/mL of mPEG-Silane (30kDa, PSB-2014, Creative PEG works), 96% ethanol and 0.1%(v/v) HCl. The slide glasses were then dried and stored at 4°C.

To make microwell chip on a cover glass, the PDMS stamp was placed on the cleaned cover glass (20 mm × 20 mm, No.1), facing the pillar surface of the stamp onto the glass. NOA81(Norland Products) drop was put at the side of the PDMS stamp to fill the space between the PDMS pillars with NOA81. The NOA81 was cured by exposing it to UV light (UVKUB2, 100%, 12 min). The PDMS stamp and excess NOA81 were then removed.

### Sample preparation

To make a reaction chamber, a cover glass with microwell chip was first oxidized in plasma cleaner (diener) for 2 min at power of 80%. The cover glass with microwell chip was attached onto the silane-PEG coated slide glass with two double-sided tapes (70 μm thickness), facing the side of microwell chip to the slide glass. The SUV solution (0.5 mg/mL) was introduced into the chamber and incubated for 10 min to make a supported lipid bilayer on the surface of microwell chip. It was washed with the SUV buffer to remove excess SUV and then washed with HKEM buffer supplemented with 0.1% BSA. Unless otherwise noted, microtubule and actin assembly were induced by diluting tubulin dimers (20% labelled) and/or actin monomers (10% labelled) in the reaction mixture containing the TicTac buffer (10 mM Hepes, 16 mM Pipes (pH 6.8), 50 mM KCl, 1 mM EGTA, 5 mM MgCl_2_) supplemented with 5% BSA, 4.4 mM GTP, 2.7 mM ATP, 20 mM DTT, 20 μg/mL catalase, 3 mg/mL glucose, 100 μg/mL glucose oxidase. Microtubule aster formation was induced by adding microtubule seeds coated beads (aMTOCs) into the mixture.

To induce branched actin assembly from the edge of microwells, the SUV solution containing DSPE-PEG-Biotin (Type 2 SUV) was used for lipid coating. Before introducing the reaction mixture into the chamber, HKEM buffer containing 200 nM of snap-streptavidin-WA (used as an NPF) and 0.1% BSA was loaded into the chamber and incubated for 5 min. The excess WA was then removed by perfusing HKEM buffer supplemented with 0.1% BSA. 80 nM of the Arp2/3 complex was added in the reaction mixture. To make asymmetric cortical actin structures, 100 nM of GST-α-actinin 4 was also added.

The reaction mixture was introduced into the chamber immediately after the preparation of the mixture. Mineral oil (Paragon Scientific, RTM13) was then loaded into the chamber in order to close the wells. Unless otherwise noted, the chamber was incubated at room temperature (22-23°C) in order to prevent tubulin precipitation. The final position of the aMTOCs was analyzed at 2 hours after sample preparation. Experiments were repeated to confirm the reproducibility.

For initial screening of biochemical conditions (in Figure 1C and Supplementary Figure 2), TicTac buffer supplemented with 0.1% BSA, 1 mM GTP, 2.7 mM ATP, 20 mM DTT, 20 μg/mL catalase, 3 mg/mL glucose, 100 μg/mL glucose oxidase was used as a control buffer solution. BSA (Sigma, A7030), Polyethylene Glycol (PEG) 3k (Sigma, P3640), PEG 20k (Sigma, 95172), Glycerol (CARLO ERBA), Ficoll400 (Sigma, F9378) and Dextran 40k (Sigma, 31389) were added in the solution at the indicated concentrations. The samples were incubated at the indicated temperatures.

### Microscopy

Microtubule asters and actin filaments in microwells were visualized using a confocal spinning-disc system (EclipseTi-E Nikon inverted microscope equipped with a CSUX1-A1 Yokogawa confocal head, an Evolve EMCCD camera (Photometrics), a Nikon CFI Plan-Apo ×60 NA1.4 oil immersion objective, a Nikon CFI S-Fluor ×100 NA1.30 oil immersion objective and a Nikon x20 NA0.75 dry objective). Time-lapse imaging was performed using Metamorph software (Universal Imaging). For time-lapse imaging of microtubule asters in microwells, images were taken every 5 or 10 min to avoid photo-damages of microtubules. Photo-bleaching experiments of lipids were also performed on this system using an iLas^2^ device.

To measure the position of aMTOC in microwells, the images were acquired using an upright Axioimager M2 Zeiss microscope equipped with an EC Plan-Neofluar dry ×20 NA0.5 dry objective and CoolSNAP EZ (Photometrics) or using a confocal spinning-disk system.

To visualize individual microtubules and actin filaments, an objective-based total internal reflection fluorescence (TIRF) microscopy instrument composed of a Nikon Eclipse Ti, an azimuthal iLas^2^ TIRF illuminator (Roper Scientific), a ×60 NA1.49 TIRF objective lens, a x100 NA1.49 TIRF objective lens and an Evolve EMCCD camera (Photometrics) was used. This system was also used to visualize tubulin precipitation in microwells. Excitation was achieved using 491, 561 and 642 nm lasers.

### Image processing and measurements

To visualize microtubule asters in microwells, the images were processed to improve the signal-to-noise ratio. Background subtraction was performed using Fiji (NIH). To further improve the signal-to-noise ratio, deconvolution was also performed for some of the images (Figure 1E, 3D and 4D) using DeconvolutionLab2 in Fiji before background subtraction. Background subtraction was not performed to show free tubulin signals in Figure 1C and Supplementary Figure 2A-E. Maximum projection was performed to show microtubule asters using Fiji. To visualize actin networks along the vertical edges, maximum projection was performed excluding the bottom and the top images of microwells (partial maximum projection). Microtubule length was manually measured using a 3D distance measurement tool in Fiji. Kymograph and temporal-color coded images were generated using plugins in Fiji.

Measurement of the center (centroid) of wells and the center of aMTOCs was performed using Fiji with bright-field images. Microwells containing only a single aMTOC were analyzed. Distance from the aMTOC to center of the well was measured from each XY coordinates. The actin inner zone (AIZ) was determined by setting thresholds of the fluorescence signals of actin. Then, the center of the AIZ was measured. The angles of the aMTOC and the center of the AIZ relative to the well center were calculated from each XY coordinate. To analyze aMTOC positioning relative to the center of the AIZ, wells were reoriented based on each XY coordinates in order to align the angles from the well center to center of the AIZ (See also Supplementary Figure 8C-D).

### Numerical simulations

Simulations were performed using Cytosim software ^57^. The motion of elastic filaments and solids surrounded by a viscous fluid was calculated using Langevin dynamics ^57^. The main parameters used in the simulation were presented in Supplementary Table 1.

In the simulations, repulsive steric effects between actin filaments and microtubule aster were considered. As a limitation in the simulation, steric repulsion also occurs between actin filaments and between microtubules at the same steric force. Attractive steric forces between filaments were not included in the simulations. The steric parameters were adapted from the range in previous studies testing steric interactions between microtubules or actin filaments using Cytosim ^58,59^. Because of excessive computational costs, it was difficult to perform simulations of centrosome positioning with dense actin filaments. Therefore, to reduce computational costs, the effective diameter of microtubules and actin filaments was set to 100 nm ^58,59^ and the cell size was set to 10 μm in radius. In addition, the total time simulated was set to 150 to 250 seconds. The simulations were performed in the two-dimensional mode.

Bulk actin network was made by adding actin filaments in the cell without fixation of their position. To make actin meshwork near the cell periphery, the actin nucleation factors and branching factors were positioned near the cell periphery (within 7 to 9.2 μm from the cell center). The position of actin nucleation factors was fixed, so that the position of one of the ends of actin filaments was fixed at the initial position. When the actin branching factor binds to an existing actin filament, it nucleates a new actin filament from the existing filament. Asymmetric actin network was made by asymmetrically localizing the actin nucleation factors and the actin branching factors.

## Statistics

Statistical tests were performed using R statistical software. Statistical test, sample sizes and P values are described in each figure legend.

**Supplementary Figure 1.**
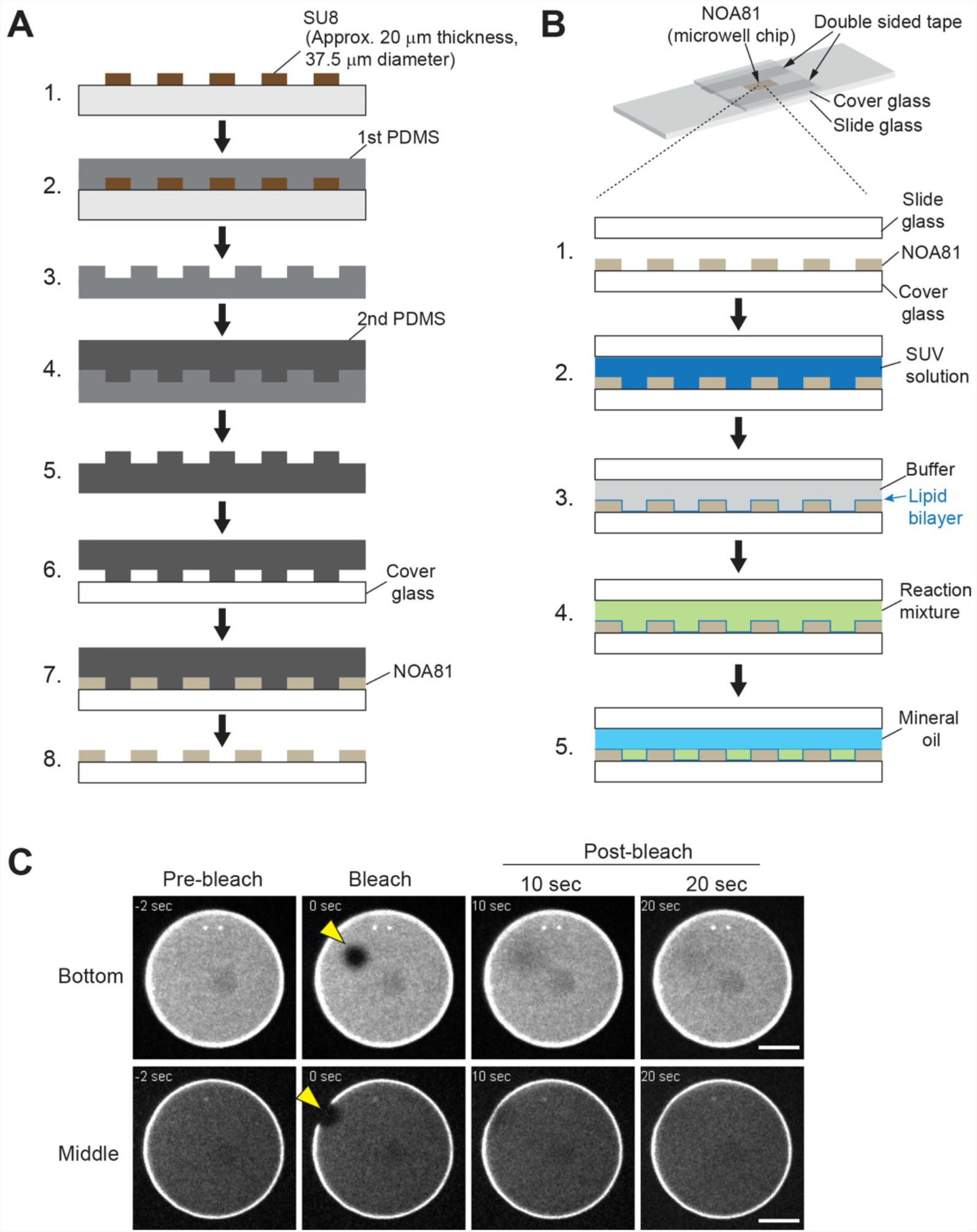
Preparation of microwells and samples. **(A)** Scheme of construction of NOA81-based microwells on a cover glass (See also Methods). SU8 mold was made on the wafer and silanized. The mold was used to make 1^st^ PDMS mold. The 1^st^ PDMS was silanized and then used to make 2^nd^ PDMS. The 2^nd^ PDMS was cut to small pieces and used as PDMS stamps. The PDMS stamp was placed on a cleaned cover glass. The space between pillars were filled with NOA81. By UV exposure, NOA81 was cured. The PDMS stamp and the excess NOA81 were removed. **(B)** Scheme of sample preparation (See also Methods). The NOA81-attached cover glass was exposed to plasma. After plasma treatment, the NOA81-attached cover glass was attached onto a silane-PEG coated slide glass with double-sided tapes. Then, the SUV solution was loaded into the chamber and incubated to make a supported lipid bilayer on the surface of microwells. The excess SUVs were then removed from the chamber by perfusing with buffer solution. After washing, the chamber was filled with the reaction mixture containing tubulin and actin. Then, the microwells were immediately closed with mineral oil. **(C)** Photo-bleaching of lipids on microwell (Bottom or middle edge of the well). Fluorescently labelled lipids were photo-bleached (shown with yellow arrow head). Fluorescent recovery indicates diffusion of lipids. Scale bar 10 μm.

**Supplementary Figure 2.**
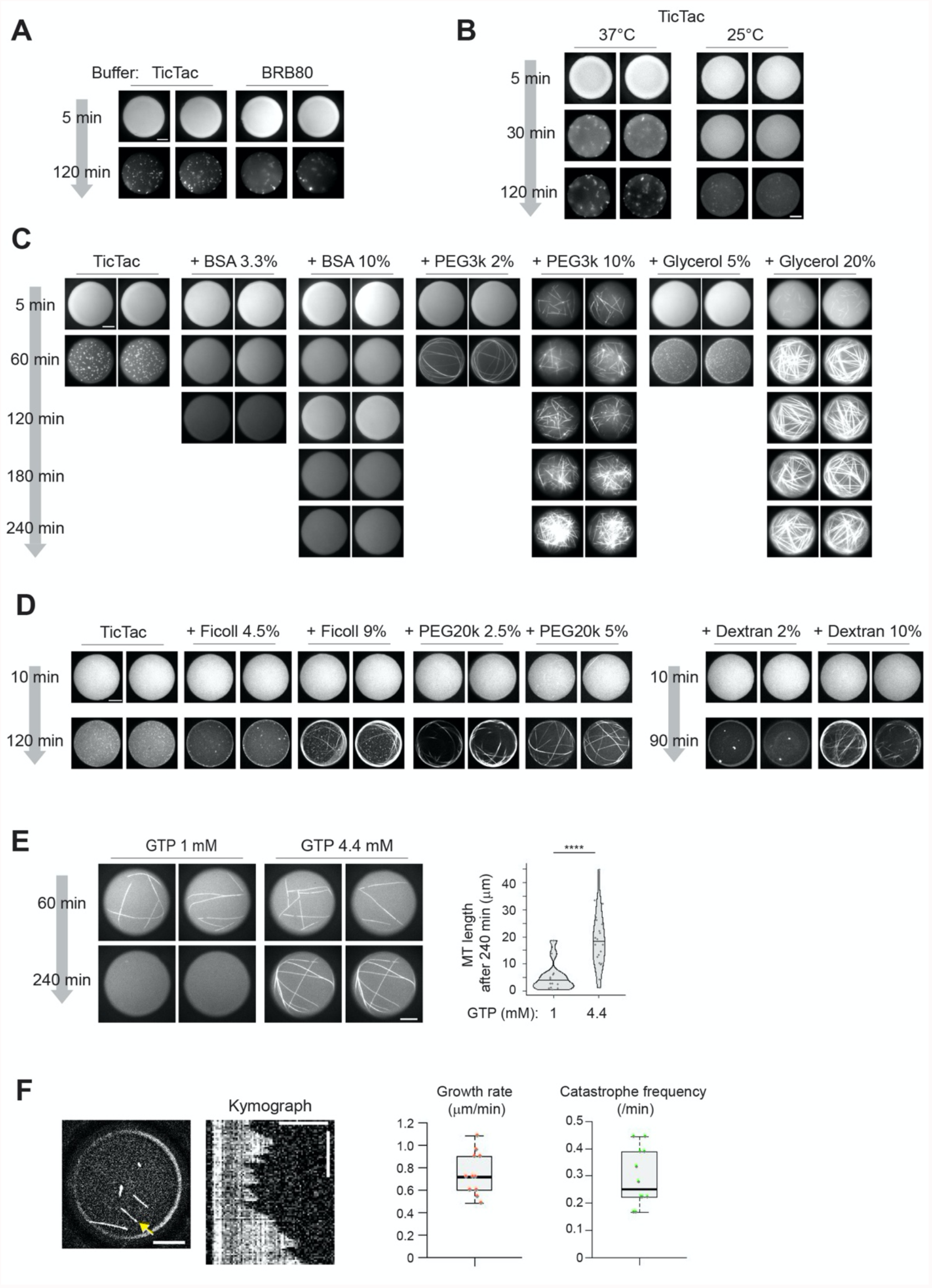
Screening of biochemical conditions to slow down tubulin precipitation. **(A)** Comparison of two different buffer solutions. Images show fluorescently labeled tubulin in microwells. Samples were incubated at 37°C after sample preparation. Tubulin 15 μM. Tubulin precipitation occurred both in the BRB80 and the TicTac buffer. **(B)** Comparison of two different temperatures in the TicTac buffer. Images show fluorescently labeled tubulin in microwells. After sample preparation, samples were incubated at the indicated temperatures. Tubulin 20 μM. Incubation at lower temperature was better to slow down tubulin precipitation, although even at the lower temperature, tubulin precipitation occurred before 2 hours after sample preparation. **(C)** and **(D)** Test of various crowding reagents. Indicated reagents were added in the TicTac buffer. Images show fluorescently labeled tubulin in microwells. Samples were incubated at 25°C. Tubulin 12.5 μM. High concentration of BSA slowed down tubulin precipitation. Some of the crowding reagents (e.g. PEG) induced MT nucleation. PEG generated tubulin aggregates, which resulted in the formation of aster-like structures. **(E)** Comparison of GTP concentrations. MT formation was induced by adding GMPCPP-stabilized MT seeds. Tubulin 8 μM. (GTP 1 mM n= 13, GTP 4.4 mM n = 20 MTs). Violin plots were shown with the median (horizontal line). ****p<0.0001 (Mann-Whitney U test). In this experiment, 15% BSA was added in the TicTac buffer. Addition of higher concentration of GTP maintained MTs for a long time. **(A)**-**(E)** Images of wells were randomly taken at each time point, indicating that the represented wells were not identical through time points. In these experiments, microwells were coated with Silane-PEG30k. As a basic buffer solution (control), BRB80 or TicTac buffer supplemented with 0.1% BSA, 1 mM GTP, 2.7 mM ATP, 20 mM DTT, 20 μg/mL catalase, 3 mg/mL glucose, 100 μg/mL glucose oxidase was used. Scale bar 10 μm. **(F)** MT dynamics in lipid coated microwells. Kymograph of the MT indicated with yellow arrow was shown. GMPCPP-stabilized MT seeds were added to induce MT formation. Tubulin 16 μM. TicTac buffer supplemented with 5% BSA, 4.4 mM GTP, 2.7 mM ATP, 10 mM DTT, 20 μg/mL catalase, 3 mg/mL glucose, 100 μg/mL glucose oxidase. Samples were incubated at 22°C. In (F), background subtraction was performed to increase signal-to-noise ratio. Scale bar 10 μm in larger image and 5 μm in kymograph, Time scale bar indicates 10 min in kymograph. n = 12 MTs. Plots show box (25 to 75%), whisker (10 to 90%). Lines in the box indicate medians.

**Supplementary Figure 3.**
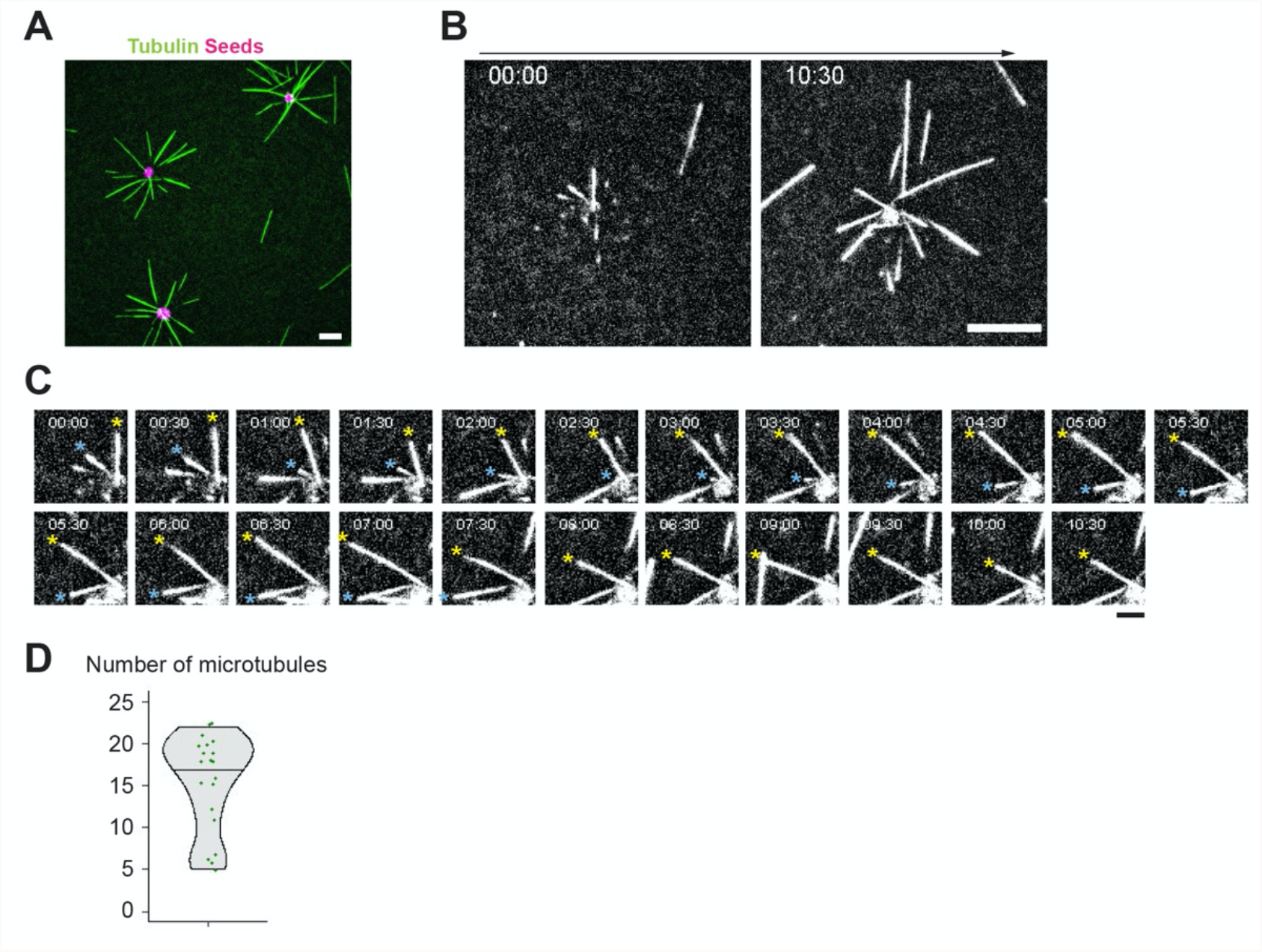
Characterization of an artificial MTOC. **(A)** Observation of MT aster formation using TIRF microscope. Scale bar 10 μm. **(B)** Time-lapse imaging of the MT aster formation using TIRF. Scale bar 10 μm. **(C)** Magnified images of (B). Time-interval 30 sec. Time bar indicates (min:sec). MTs showing dynamic instability were indicated with asterisks. Scale bar 5 μm. **(D)** Number of MTs emanating from the aMTOCs in microwells. Tubulin 18 μM. n = 20 wells.

**Supplementary Figure 4.**
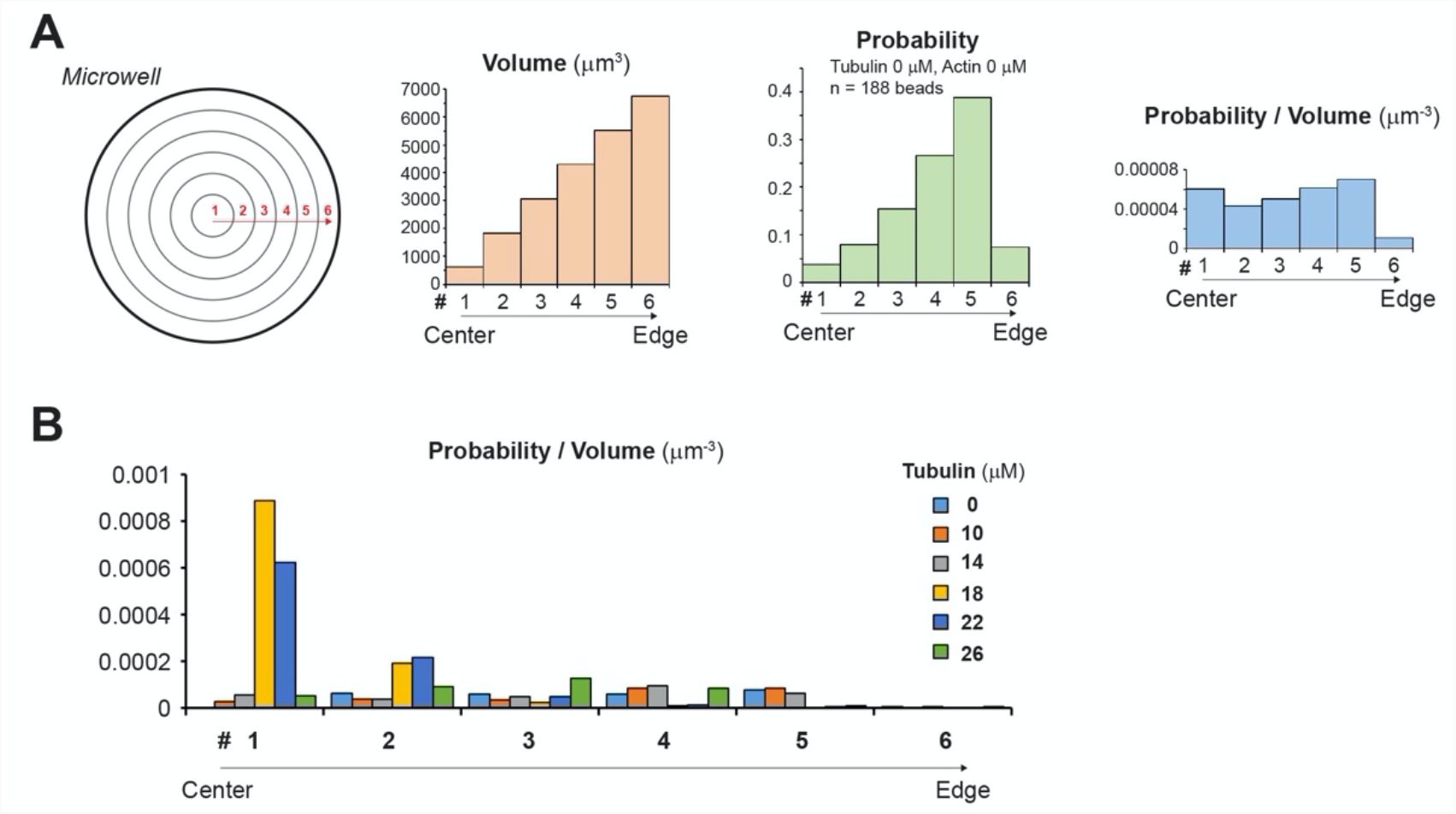
Characterization of aMTOC positioning in microwells. **(A)** Consideration of the probability of aMTOC distribution. When the volumes of 6 segments (equal intervals from well center) are considered as shown in the left scheme, the volume per segment increases from well center toward well edge (left histogram), suggesting that probability of the aMTOC distribution increases from well center toward the edge. Middle histogram shows experimental probability of the aMTOC position in the absence of free tubulin and actin (n = 188 wells). The probability tended to increase from well center toward the edge. However, the distribution near the edge was restricted, because of the size of aMTOCs (bead 1.5 μm radius + MT seeds). Right histogram indicates probability per volume, suggesting almost random distribution of the aMTOCs in microwells. Volume per segment was calculated based on the approximate size of microwells (37.5 μm in diameter and 20 μm in height). **(B)** Distribution of aMTOC in microwells at the indicated tubulin concentrations. Probability per volume was calculated as shown in (C). Data shown in Figure 1F were used.

**Supplementary Figure 5.**
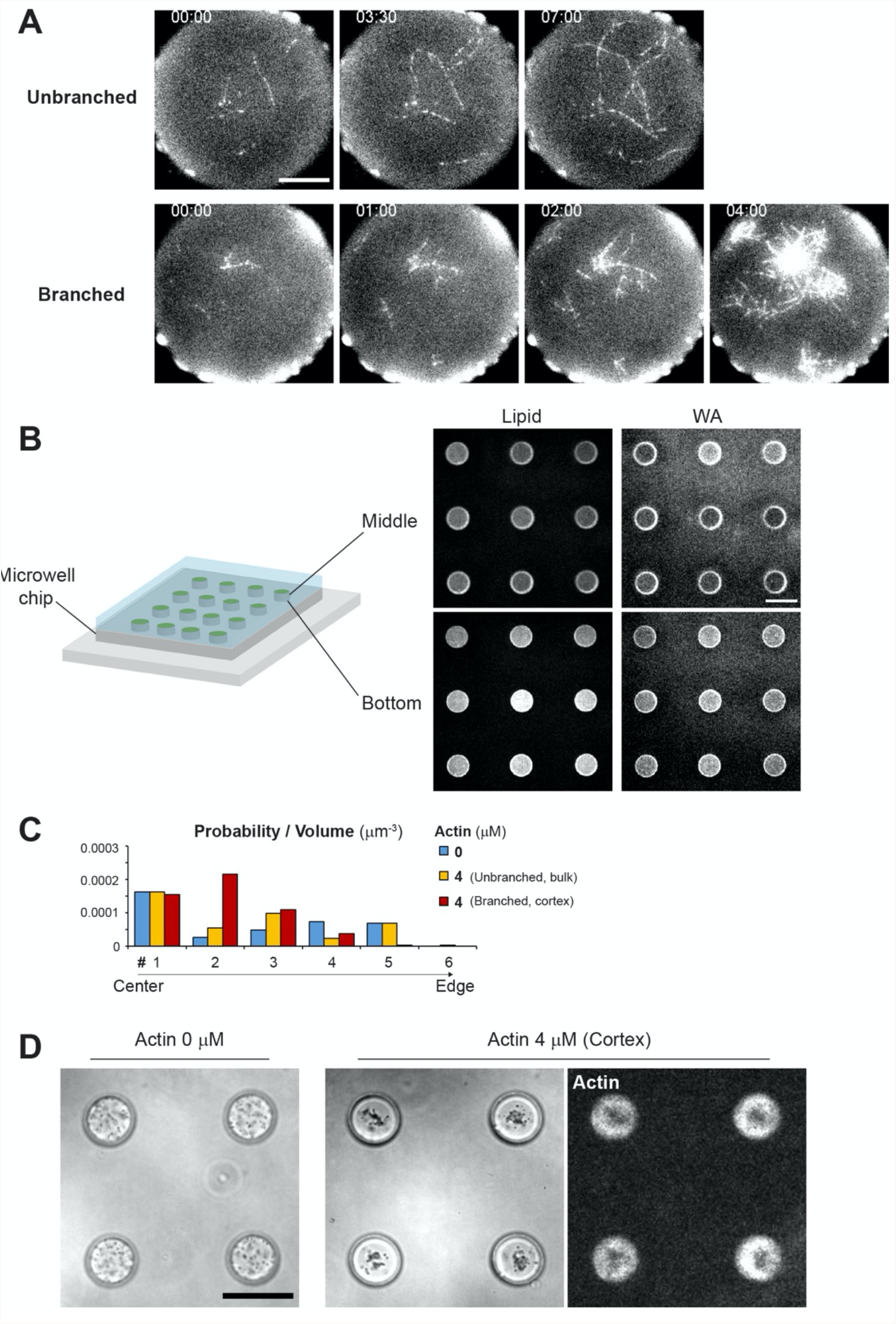
Characterization of actin assembly in microwells. **(A)** Time-lapse imaging of actin assembly on lipid coated microwells by TIRF microscope. Top images show unbranched actin (Actin 1.25 μM). Bottom images show branched actin (Actin 1.25 μM, Arp2/3 complex 80 nM, GST-WA 100 nM). In these experiments, 0.25% of methyl cellulose (Sigma, 1500 cP) was added to visualize actin filaments within the TIRF field. Time indicates (min:sec). Scale bar 10 μm. **(B)** Lipid and NPF (WA) coated microwells. Fluorescence labelled lipid and snap-streptavidin-WA were used. Scale bar 50 μm. **(C)** Distribution of the aMTOC in the absence of free tubulin in microwells. Probability per volume was calculated as shown in Supplementary Figure S4(A). Data shown in Figure 2E were used. **(D)** Distribution of smaller beads (1 μm in diameter, PolySciences, #08226) in the absence or presence of cortical actin (Actin 4 μM, Arp2/3 80 nM and NPF coating) in microwells. In bright field images, black dots in microwells indicate the beads. The presence of cortical actin clustered the beads to the well center. Scale bar, 50 μm.

**Supplementary Figure 6.**
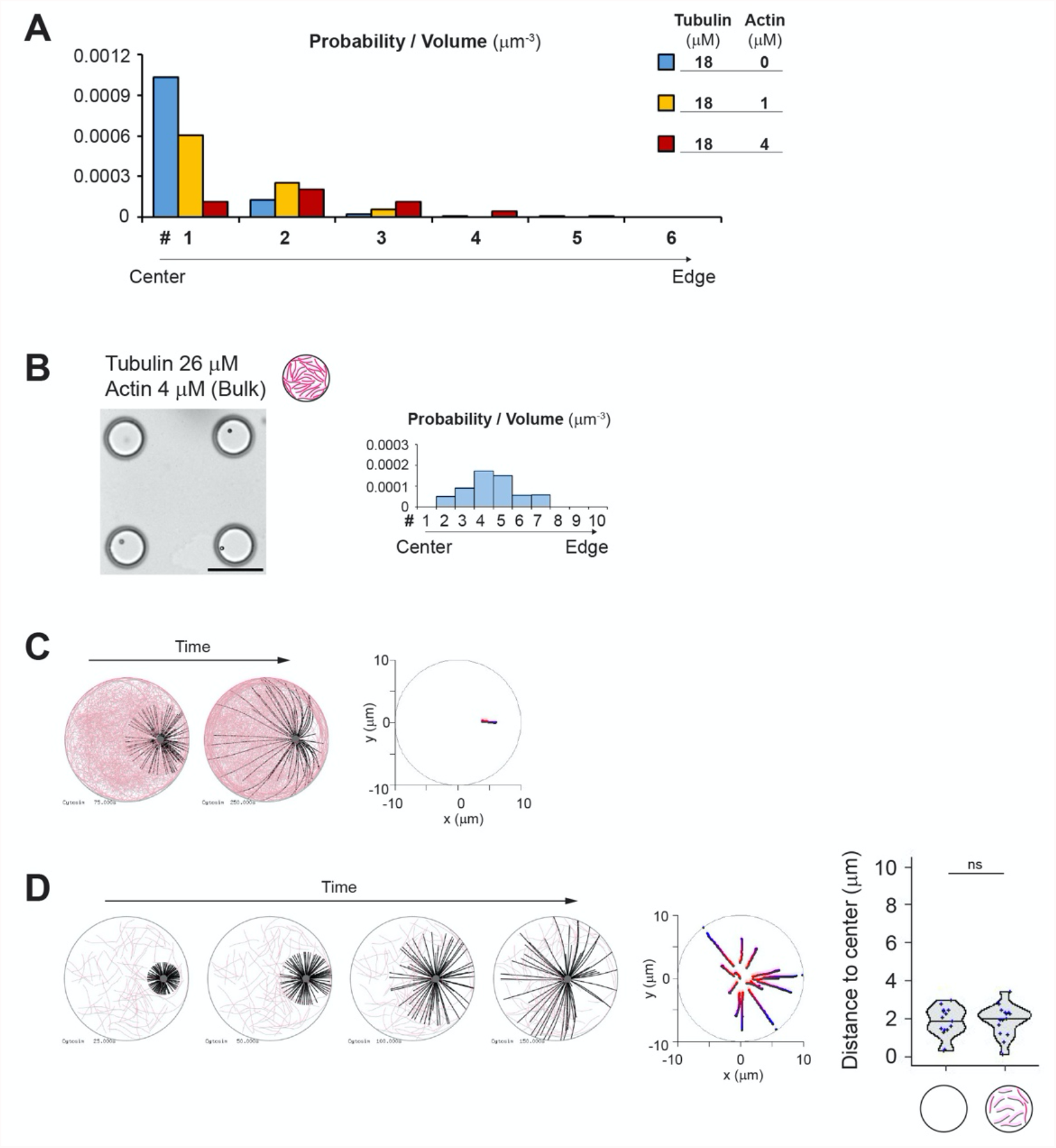
aMTOC positioning in the presence of bulk actin network. **(A)** Distribution of aMTOC in microwells in the presence of the indicated tubulin and actin concentrations. Probability per volume was calculated as shown in Supplementary Figure 4A. Data shown in Figure 3B were used. **(B)** Distribution of the aMTOC in microwells in the presence of tubulin 26 μM and bulk actin 4 μM. Probability per volume was calculated as shown in Supplementary Figure 4A. Data from Figure 3B. Scale bar, 50 μm. **(C)** Simulation in the presence of bulk actin network. Even with longer MT formation (compared to Figure 3E), the MTOC centering was not occurred. Different time points (75 and 250 sec) were shown. Right graph shows trajectories from blue (0 sec) to red (250 sec). MTOC, gray, MT, black, Actin, pink. **(D)** Simulations in the presence of lower density of actin. Different time points (25, 50, 100 and 150 sec) were shown. Middle graph shows trajectories from blue (0 sec) to red (150 sec). Right graph shows final position of MTOC (at 150 sec). Data of the absence of actin (left) are same as shown in Figure 3G. ns (not significant)>0.1 (Mann-Whitney U test). 15 simulations per condition. Violin plots were shown with the median (horizontal line).

**Supplementary Figure 7.**
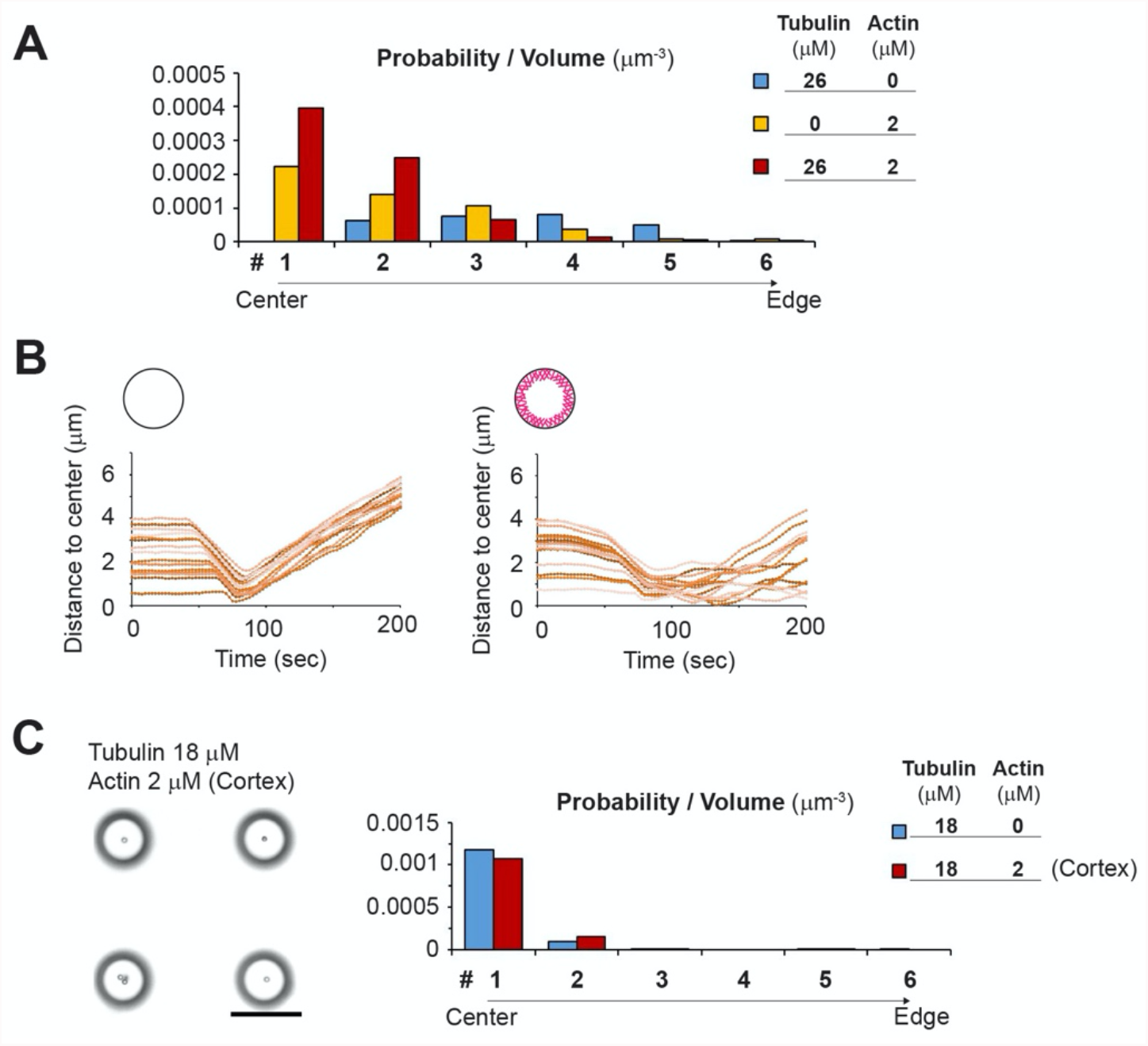
aMTOC positioning in the presence of cortical actin network. **(A)** Distribution of the aMTOC in microwells in the presence of the indicated tubulin and actin concentrations. Probability per volume was calculated as shown in Supplementary Figure 4A. Data shown in Figure 4B were used. **(B)** Simulations of MTOC position over time in the absence (left) or presence (right) of cortical actin. 15 simulations per condition were shown with different colors. **(C)** Distribution of the aMTOC in microwells in the presence of tubulin 18 μM and cortical actin 2 μM. Probability per volume was calculated as shown in Supplementary Figure 4A. Data shown in Figure 4B were used. Scale bar, 50 μm.

**Supplementary Figure 8.**
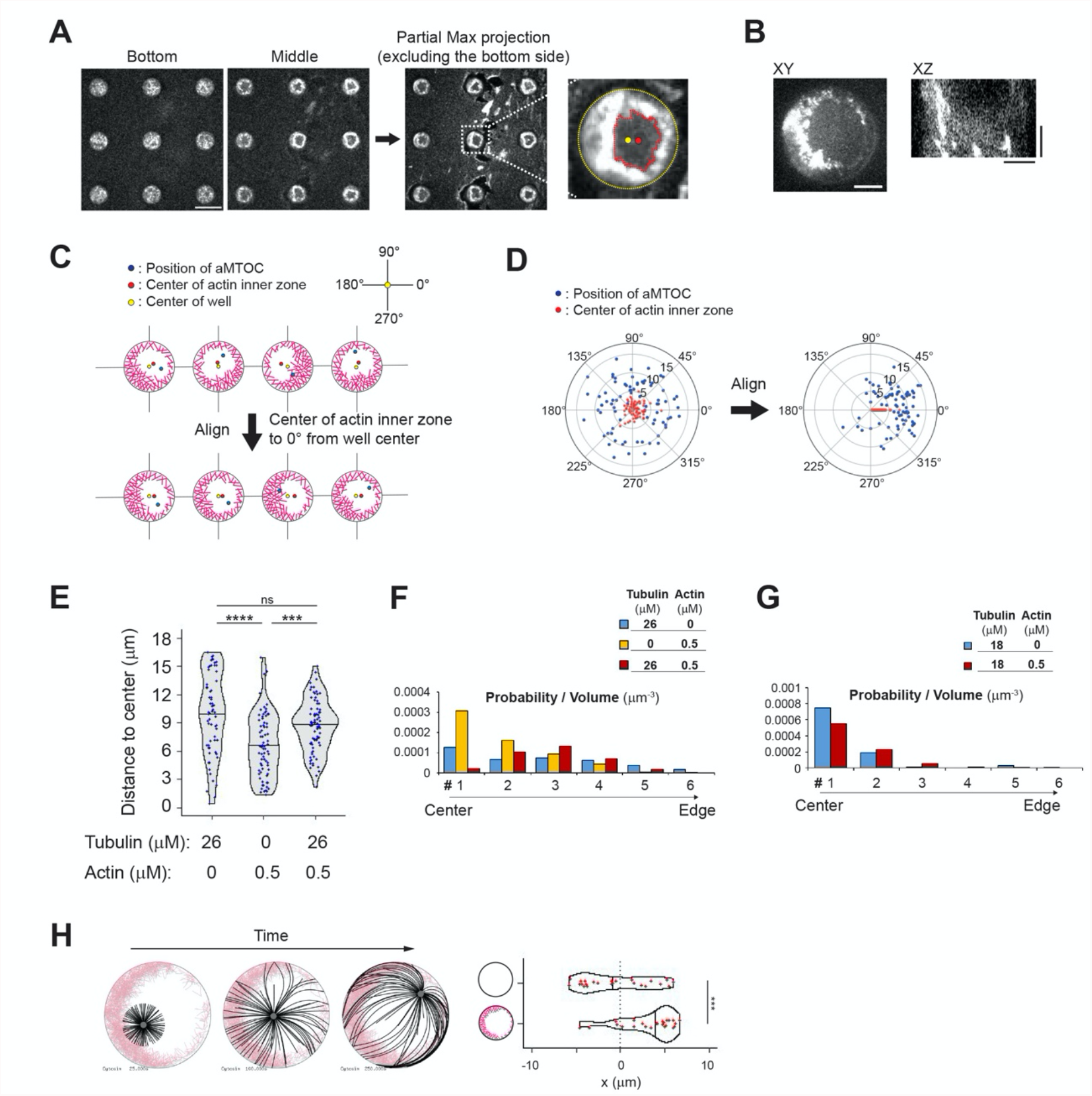
aMTOC positioning in the presence of asymmetric actin network. **(A)** Detection of actin inner zone. Actin 0.5 μM and a-actinin 100 nM with Arp2/3 complex and NPF (WA) coating. The bottom and top were excluded from the maximum projection to visualize the vertical edges of actin. Using the partial max projection images and defining a threshold of actin intensity, the region of actin inner zone was detected and the center (centroid) was measured. Scale bar, 50 μm. **(B)** Representative image of actin shown in Figure 5H (120 min). XY and XZ view of the well were shown. Scale bar, 10 μm. The actin layer was present throughout the lateral edge of microwell. **(C)** and **(D)** To analyze the aMTOC position with respect to the asymmetry of cortical actin, the angles from the well center to the center of the actin inner zone were aligned at the same degree (0°). It means that wells were reoriented in order to align the angles from the well center to the center of the actin inner zone. **(E)** Distance from aMTOC to well center (2 hours after sample preparation). Data shown in Figure 5E, F and I were used. (Tubulin 26 μM Actin 0 μM n = 65, Tubulin 0 μM Actin 0.5 μM n = 74, Tubulin 26 μM Actin 0.5 μM n = 79 wells) ***p<0.001, ****p<0.0001, ns (not significant)>0.1 (Kruskal-Wallis test with Dunn’s multiple comparison test). **(F)** and **(G)** Distribution of the aMTOC in microwells in the presence of tubulin and actin at the indicated concentrations. Probability per volume was calculated as shown in Supplementary Figure 4A. Data shown in Supplementary Figure 8E and Figure 5J were used for (F) and (G), respectively. **(H)** Simulations in the presence of asymmetric actin when the initial position was randomly chosen within 4 μm from cell center. 25 simulations per condition. Different time points (From left, 25, 100 and 250 sec) were shown. Even if the initial position is off-centered, the MTOC tended to migrate toward the thinner side of the actin network. ***p<0.001 (Mann-Whitney U test). Violin plots were shown with the median (horizontal line).

**Supplementary Movie 1. Centering of MT aster in the absence of actin**.

Time-lapse imaging of MT aster formation at 18 μM of tubulin in the absence of actin. Data is also shown in Figure 1G. Scale bar, 10 μm.

**Supplementary Movie 2. Off-centering of MT aster in the absence of actin**.

Time-lapse imaging of MT aster formation at 26 μM of tubulin in the absence of actin. Data is also shown in Figure 1I. Scale bar, 10 μm.

**Supplementary Movie 3. Simulation of MT aster positioning in the absence or presence of bulk actin filaments**.

Simulations in the absence (left) or presence (right) of bulk actin filaments. From 25 to 150 sec. MTOC, gray, MT, black, Actin, pink. Data is also shown in Figure 3E.

**Supplementary Movie 4. MT aster centering in the presence of cortical actin network**. Time-lapse imaging of MT aster formation at 26 μM of tubulin in the absence (left) or presence (right) of cortical actin. Data is also shown in Figure 4E. Scale bar, 10 μm.

**Supplementary Movie 5. Simulation of MT aster positioning in the absence or presence of actin filaments near cell periphery**.

Simulations in the absence (left) or presence (right) of actin filaments near cell periphery. From 25 to 200 sec. MTOC, gray, MT, black, Actin, pink. Data is also shown in Figure 4I.

**Supplementary Movie 6. MT aster positioning in the presence of asymmetric actin network**. Time-lapse imaging of MT aster formation at 26 μM of tubulin in the presence of asymmetric actin network. Data is also shown in Figure 5G. Scale bar, 10 μm.

**Supplementary Movie 7. Simulation of MT aster positioning in the absence or presence of asymmetric actin network**.

Simulations in the absence (left) or presence (right) of asymmetric actin network near cell periphery. From 25 to 250 sec. MTOC, gray, MT, black, Actin, pink. Data is also shown in Figure 5K.

## Notes

### Competing Interest Statement

The authors have declared no competing interest.

