## SupplementaryTable S1 for "The architecture of the actin network can balance the pushing forces produced by growing microtubules"

### Supplementary Table 1

#### Cytosim parameters

| | | Value | Note |
|---|---|---|---|
| Global | Time step | 0.01 s | Computational parameter |
| | Viscosity | 0.3 pN s/μm$^2$ | (Ref S1) |
| | Steric force constant (Repulsion) | 1.5 pN/μm | Adapted from the range in previous studies (Ref S2, S3) |
| Cell | Radius | 10 μm | Radius for the basic circular geometry (Ref S4) |
| | Confinement stiffness | 500 pN/μm | Confinement strength of microtubules and actin filaments inside the cell (Ref S4) |
| Microtubule | Rigidity | 25 pN μm$^2$ | Persistence length Lp = 5200 μm (Red S4, S5) |
| | Segmentation | 0.2 μm | Computational parameter |
| | Steric radius | 50 nm | (Ref S2, S3) |
| | Growing speed | Varied | (Ref S4, S6) |
| | Stall force | 1.67 pN | Growing sensitivity to force (Ref S7, S8) |
| | Shrinking speed | 0.27 μm/s | (Ref S4, S6) |
| | Catastrophe rate | 0.01, 0.04 s$^{-1}$ | Unloaded and stalled catastrophe rate (Ref S4, S9) |
| | Rescue rate | 0.064 s$^{-1}$ | (Ref S4, S6) |
| | Number of microtubules | 90 | Number of microtubules at t=0 sec |
| | Initial length | 1 μm | Length of microtubules at t=0 sec |
| MTOC | Radius | 0.5 μm | Radius of MTOC (Ref S4) |
| | First anchoring stiffness | 500 pN/μm | Stiffness of the link anchoring microtubules to the center of MTOC (Ref S4) |
| | Second anchoring stiffness | 500 pN/μm | Stiffness of the link anchoring microtubules to a point on the MTOC periphery (Ref S4) |
| Actin | Rigidity | 0.06 pN μm$^2$ | Persistence length Lp = 15 μm was chosen. (Ref S2, S10) |
| | Segmentation | 0.2 μm | Computational parameter |
| | Steric radius | 50 nm | (Ref S2, S3) |

Bulk actin network

| | | | |
|---|---|---|---|
| Actin | Number | Varied | Number of actin filaments |
| | Length | 5 μm | Length of actin filaments |

Actin near cell periphery

| | | | |
|---|---|---|---|
| Actin nucleation factor | Nucleation rate | 100 $s^{-1}$ | Rate of nucleation, High enough for quick actin assembly |
| | Length of actin filaments | 2 μm | |
| | Unbinding rate | 0 $s^{-1}$ | No detachment |
| | Stiffness | 100 pN/μm | Stiffness of the link between the nucleator and its fixed anchoring position |
| | Number | Varied | Number of actin nucleation factor |
| Actin branching factor | Nucleation rate | 100 $s^{-1}$ | Nucleation rate when bound to an existing filament, for quick actin assembly |
| | Length of actin filaments | 1 μm | |
| | Binding rate | 1 $s^{-1}$ | Binding rate for quick actin assembly |
| | Binding range | 0.01 μm | Bind to a close filament |
| | Unbinding rate | 0 $s^{-1}$ | No detachment |
| | Equilibrium angle | 1.22 rad | Angle between the two branches, 70° (Ref S11) |
| | Angular stiffness | 0.13 pN.μm/rad | Stiffness of the torque connecting the two branches (Ref S2, S11) |
| | Diffusion | 0 $\mu m^2/s$ | No diffusion, in order to limit the region of actin assembly |
| | Number | Varied | Number of actin branching factor |

**Varying parameters**

| | | Value | Figures | Note |
|---|---|---|---|---|
| Microtubule | Growing speed | 0.07 μm/s | Fig.3E-G Sup.Fig.6C-D | Aster with shorter MTs |
| | | 0.13 μm/s | Fig.4I-K, 5K-M Sup.Fig.7B, 8H | (Ref S4, S6) |
| Actin (Bulk actin) | Number | 800 | Fig.3E-G, Sup.Fig.6C | Dense actin filaments (Bulk) |
| | | 80 | Sup.Fig.6D | Loose actin filaments (Buk) |
| Actin nucleation factor | Number | 1400 | Fig.4I-K Sup.Fig.7B | Symmetric actin network (periphery) |
| | | 905 | Fig.5K-M, Sup.Fig.8H | Asymmetric actin network (periphery) |
| Actin branching factor | Number | 2100 | Fig.4I-K Sup.Fig.7B | Symmetric actin network (periphery) |
| | | 365 | Fig.5K-M, Sup.Fig.8H | Asymmetric actin network (periphery) |
